# Klf4 overexpression remodels chromatin to reprogram late retinal progenitors toward a retinal ganglion cell-like fate

**DOI:** 10.64898/2026.06.28.734840

**Authors:** VM Oliveira-Valença, JM Roberts, F (常菲) Chang, A Bosco, ML Vetter, MS Silveira

## Abstract

Developing neuron-replacement therapies for retinal ganglion cells (RGCs) lost to injury or disease requires a deeper understanding of how restriction to cell identity acquisition may be overcome. Previously, we showed that overexpression of *Klf4* in late retinal progenitor cells (late RPCs), which are normally restricted from RGC production, is sufficient to produce cells that display a subset of canonical RGC properties including RGC-associated gene expression and morphological features. In the present study, we investigated the transcriptional and epigenetic mechanisms by which *Klf4* overexpression influences the fate of cell types generated from late RPCs. scRNA-seq analysis revealed that *Klf4* induces transcriptional changes, with some cells exhibiting gene expression profiles similar to those of resident RGCs. In addition, we observed widespread changes in chromatin accessibility, suggesting that KLF4 remodels the chromatin of late RPCs and influences their transcriptional profile. Our findings show KLF4-driven reprogramming of late RPCs, providing insight into progenitor competence and fate specification to an RGC-like identity. These results suggest that KLF4 could be a component in regenerative therapies due to its ability to reprogram and induce RGC genes outside of the normal RGC developmental window.

## INTRODUCTION

Retinal development is orchestrated through a highly regulated sequence of events that produces all retinal neurons and Müller glia from multipotent retinal progenitor cells (RPCs). This process is directed by gene regulatory networks that are temporally coordinated to ensure proper specification and differentiation of cellular identities. The generation of these retinal cell types follows a controlled progression across two major developmental windows (Cayouette et al., 2006; Livesey & Cepko, 2001). During embryonic development, early RPCs give rise to ganglion cells, amacrine cells, horizontal cells, and cone photoreceptors. As development proceeds, epigenetic and transcriptional changes alter the potency of RPCs, allowing late RPCs in the postnatal stage to produce rod photoreceptors, Müller glia, bipolar cells, and additional amacrine subtypes (Lyu et al., 2021; Shiau et al., 2021).

A major challenge is to determine whether specific cell types can be generated outside of their restricted developmental window, with the goal of driving cell reprogramming to promote regeneration of retinal neurons lost to injury or disease. The transcription factor Krüppel-like factor 4 (KLF4) is recognized as one of the four Yamanaka factors and plays a central role in cell reprogramming (Takahashi et al., 2007; Yamanaka, 2008). It is also a potent developmental regulator that can function as both a transcriptional activator and repressor (Evans et al., 2007; Nakatake et al., 2006; Shie et al., 2000). During retinal development, KLF4 is expressed across all stages, but postnatally, its expression becomes confined to the inner retina—specifically the inner nuclear layer (INL) and ganglion cell layer (GCL) (Njaine et al., 2014). While KLF4 deletion does not significantly disrupt retinogenesis, potentially due to functional redundancy within the KLF family (Fang et al., 2016; Jiang et al., 2008; Rocha-Martins et al., 2019), gain-of-function experiments strikingly demonstrate that overexpression in late RPCs alters cell fate, generating cells with characteristics of retinal ganglion cells, termed induced RGCs (iRGCs), including enrichment for multiple RGC-associated genes (Rocha-Martins et al., 2019).

It is unclear how similar these iRGCs are to embryonically derived RGCs, and how KLF4 induces this cell fate reprogramming process in late RPCs. To further elucidate the identity of iRGCs and the molecular alterations driven by KLF4, we employed single-cell RNA sequencing (scRNA-seq). We observed changes in transcriptional profiles of late RPC-derived cells as well as shifts in retinal cell fate with *Klf4* overexpression, with induction of an RGC-like gene expression program in a subset of cells. In parallel, we analyzed chromatin accessibility through transposase-accessible chromatin sequencing (ATAC-seq) and identified pathways affected by KLF4-induced chromatin landscape changes. Our data shows that KLF4 can broadly induce chromatin remodeling, opening genomic regions proximal to upregulated genes, including RGC genes, and closing other regions. These results suggest that KLF4 acts to remodel chromatin, partially reprogramming late progenitors to generate RGC-like cells. The mechanisms underlying KLF4-mediated alterations in fate, as well as the identity of the resultant cell populations, are the central focus of this study.

## MATERIALS AND METHODS

### Experimental animals

All experiments followed international guidelines and were approved by the Ethics Committee on Animal Experimentation by the Institutional animal Care and Use Committee at the University of Utah, where mice were housed in an Association for Assessment and Accreditation of Laboratory Animal Care (AAALAC) accredited animal facility with 12h light/12h dark cycle and ad libitum access to food and water. Both male and female neonatal C57Bl6/J mice (P0 and P1, RRID:IMSR_JAX:000664) underwent *in vivo* electroporation and were analyzed 2 and 10 days later. For the EdU trace experiment, neonatal mice received subcutaneous injections of 2.5mM EdU at P0 and P1, then underwent electroporation at P1.

### Plasmids

Plasmids pUb::CST (pUb-human ubiquitin C promoter) and pUb::GFP (Matsuda & Cepko, 2004) were kindly provided by Dr. Michael A. Dyer (St. Jude Hospital, Tennessee, USA). For *Klf4* overexpression, the plasmids referenced by Rocha-Martins et al. (2019) were employed. The final concentration of each plasmid solution was 5 µg/µL for both the control group (70% pUb::CST and 30% pUb::GFP) and the KLF4 OE group (70% pUb::Klf4 and 30% pUb::GFP).

### In vivo electroporation

This procedure followed established methods (Matsuda & Cepko, 2007; Rocha-Martins et al., 2019). Briefly, P0 mice were anesthetized by hypothermia, and the plasmid solution (0.5 µL; 5.0 µg/µL with 0.1% Fast Green Dye, Cayman Chemical #14335) was injected into the subretinal space using a Hamilton syringe with a 33G blunt-end needle. Five 80V pulses (50 ms with 950 ms intervals) were delivered via forceps-type electrode (Nepagene CUY650P7) with Neurgel (Spes Medica). Pups were returned to their cages after recovering body temperature.

### Immunofluorescence and EdU

After euthanasia, eyes were harvested and immersion-fixed in 4% paraformaldehyde in phosphate-buffered saline (PBS) overnight, cryoprotected in 30% sucrose in PBS, then immersed in OCT for transverse cryosections (10 µm), and mounted on 6% silane-treated slides (Sigma #440140). Electroporated retinas underwent immunoreaction with primary antibodies against FOXP1 (Abcam #Ab16645), POU4F2 (Rockland #600-101-MJ0) and POU4F1 (Milipore #MAB1585), combined with anti-GFP (Invitrogen mAB3E6 or A11122). The EdU assay was performed according to the manufacturer’s protocol (Click-iT EdU proliferation kit for imaging Thermo Fisher #C10340).

### Image acquisition

Retina sections were imaged using a 20x/0.8 objective by confocal microscopy (Nikon A1, Eclipse Ti2). Image acquisition settings were maintained across replicate experiments, controlled by NIS-Elements (Nikon). Image processing involved maximum intensity projection (MIP) and adjustments to color and brightness in ImageJ/Fiji. Images were cropped and rotated in Adobe Photoshop and panels prepared in Adobe Illustrator.

### scRNA-seq sample preparation

Wild-type mice at P0 were electroporated using the plasmid preparations KLF4-GFP or CST-GFP that were injected into the subretinal space. Ten days later, retinas were collected for dissociation using the same protocol as described in Oliveira-Valença et al. (2025). Single, viable, GFP^+^ cells were isolated by cell sorting with BD FACSAria 4 laser instrument using 70 µm nozzle size and collected into 20 µL DMEM + 5% FBS. Multiple retinas were combined as needed to obtain 30k GFP^+^ cells per condition: control (CTR) and *Klf4* overexpression (KLF4 OE).

### scRNA-seq Library prep and Sequencing

Library generation and sequencing were performed by the Huntsman Cancer Institute High-Throughput Genomics and Bioinformatics Analysis Shared Resource. Sorted cells were prepared with the 10X Genomics Next GEM Single Cell 3’ Gene Expression Library prep v3.1 with UDI. The library was chemically denatured and applied to an Illumina NovaSeq flow cell using the NovaSeq XP workflow (20043131). Following transfer of the flow cell to an Illumina NovaSeq 6000 instrument, a 151 x 151 cycle paired end sequence run was performed using a NovaSeq 6000 S4 reagent Kit v1.5 (20028312). Sequencing was performed to a depth of at least 250M paired reads/sample.

### scRNA-seq Analysis

Fastq files were aligned to the mouse transcriptome index for mm10 and GFP using Cellranger v7.0.1 with expected cells set to 5,000 and introns included in counting. The filtered matrix output was imported to Seurat v4.3.0 and further filtered for cells with less than 6% (*Klf4* overexpression, KLF4 OE) or 5% (CTR) mitochondrial transcripts, more than 1,200 genes and fewer than 30,000 (KLF4 OE) or 25,000 (CTR) transcripts. The final number of cells for each condition were: 5,003 CTR and 6,250 *Klf4* overexpression. The control and *Klf4* overexpression datasets were merged, and the Seurat SCTransform pipeline v0.3.5 ran with all default settings. Seurat-defined unsupervised clusters were annotated with known markers for retinal cells and combined as appropriate based on those markers. Automated cell type classification was performed with Seurat v5.0.2 and Azimuth v0.5.0 using a reference index created from a published scRNA-seq developmental dataset (Clark et al., 2019). This was compared to manual annotation, and manual annotation was used for further analysis. Cluster markers and differentially expressed genes between the experimental conditions were defined using the Seurat v5.1.0 FindAllMarkers and FindMarkers functions respectively, with default settings. The default statistical text for these functions is Wilcoxon Rank Sum test with Bonferroni correction of p-values. Integration with developmental data was achieved using Clark et al. (2019). Specifically, the developmental dataset was subset for only those cells used for pseudotime analysis, as defined in the cell metadata. The miniaturized developmental dataset and the KLF4 dataset from this study were separately run through a Seurat pipeline v5.3.1 to normalize, find variable features, scale, and perform PCA dimensionality reduction. They were combined with the IntegrateLayers function using CCA integration, and the resulting data plotted by UMAP. RGC-enriched genes were identified from the Clark et al. (2019) dataset using Seurat FindAllMarkers. The most enriched markers were defined as those with adjusted p-value < 0.05 and ordered by greatest average log2 Fold Change.

### ATAC-seq sample preparation

Retinas were electroporated at P0 with *Klf4* or CTR and dissociated 2 or 10 days later using papain. Approximately 80-100k GFP^+^ cells were sorted from each condition, for each timepoint, and ATAC-seq was performed with the Active Motif ATAC-seq kit (#105310). Library fragment length was verified with an Agilent D1000 DNA ScreenTape Assay (Agilent 2200 TapeStation). Sequencing libraries were chemically denatured and transferred to an Illumina NovaSeq X instrument. A 151 x 151 cycle paired-end sequence run was performed using a NovaSeq X Series 10B Reagent Kit (20085594). Samples were sequenced to a total depth of 52-290 million paired reads per sample. We generated a single replicate of each condition 2 days after electroporation, and 2 replicates from control and 3 replicates from *Klf4* overexpression 10 days after electroporation.

Adapters were removed with cutadapt v3.5 specifying: NextSeq-specific quality trimming with a quality threshold of 20, trimming requires that at least one base of the adapter be aligned to the read, and to discard reads shorter than 1bp. Trimmed reads were aligned to mouse genome mm39 with bowtie2 v2.2.9 using samtools v1.16 very sensitive local alignment settings, and suppressing unpaired or discordant alignments. Using samtools and bedtools v2.28.0, bam alignment files were marked for optical duplicates with distance 2,500, then filtered to remove duplicate, unaligned, secondarily aligned, mitochondrial reads, and those in blacklisted regions. Bam files from the same sample, but distinct sequencing runs were merged.

Using macs2 v2.2.7.1, we performed peak-calling on each fragment, rather than read pairs. To center peaks over the ATAC-seq cut-sites, reads were shifted 100 bp in the 5’ direction and extended 200 bp 3’ from the 5’ end. Further peak-calling parameters specified to keep all potential duplicates, used effective genome size 2.5e9, minimum peak size of 100 bp, and minimum q-value of 0.01 for peak detection. A consensus peak list was created from peak regions that were called in at least two samples. Peaks separated by less than 100bp were combined.

Quantification of reads in consensus peaks was performed by featureCounts v2.1.1 using the paired-end parameter. Peaks on contiguous chromosomes and greater than 10 counts in at least two samples were passed to DESeq2 v1.50.2 for pairwise differential accessibility analysis. Annotation was performed with ChIPseeker v1.46.1 using GenomicRanges v 1.62.1, genomic coordinates database: TxDb.Mmusculus.UCSC.mm39.knownGene v3.22.0, and gene metadata database: org.Mm.eg.db v3.22.0.

KLF4 motifs across peaks were identified with chromVAR v1.32.0 and motifmatchr v1.32.0. GC bias was added to the DESeq data set object before matchMotifs was run using KLF4 motif MA0039.2 (JASPAR) and reference genome BSgenome.Mmusculus.UCSC.mm39 v1.4.3. Gene set overrepresentation analysis was performed with the msigdbr v25.1.1 mouse gene collection, GO Biological Process, against peak gene annotation. The fgsea v1.36.0 fora function was run using peaks with increased accessibility as the query, and all peaks as the universe. Peaks annotated as distal intergenic were excluded from both lists. Plots were made with ggplot2 v4.0.2, UpSetR v1.4.0, and igraph v2.2.1.

To generate log2 fold-enrichment tracks for visualization, fragment coverage files were created from filtered, merged alignment files with Biotoolbox v2.041 bam2wig using reads per million scaling, and recoding across the entire aligned reads. Global geometric mean fragments were calculated from these coverage files for each experimental condition with MultiRepChIPSeq generate_mean_bedGraph v3.3 using genome size 2.5e9. Mac2 bdgcmp calculated the fold enrichment of fragment coverage files over the global mean. These were converted to log2-transformed bigwigs with Biotoolbox manipulate_wig, and replicates were averaged by condition with deeptools v3.5.4 bigwigAverage. Heatmap data was computed and plotted with deeptools.

Motif enrichment was performed with HOMER v5.1 using genome mm39 and size as given. Known results from HOMER were plotted with motifStack v1.54.0.

## RESULTS

### Klf4 overexpression in late retinal progenitors results in cell fate changes

We have shown previously that overexpression of *Klf4* in the late-stage retinal progenitor population alters cell fate decisions in the rat retina (Rocha-Martins et al., 2019). Here, we sought to validate these results in the mouse retina and to explore the effect of *Klf4* at the transcriptional level. To deliver genes to late RPCs, we performed *in vivo* electroporation of a *Gfp* expression vector along with *Klf4* or an empty vector as control. These two plasmids were injected into the subretinal space on the day of birth or the following day (postnatal days 0-1, P0-P1), and retinas were examined 2 or 10 days later (**Figure 1A**).

**Figure 1:**
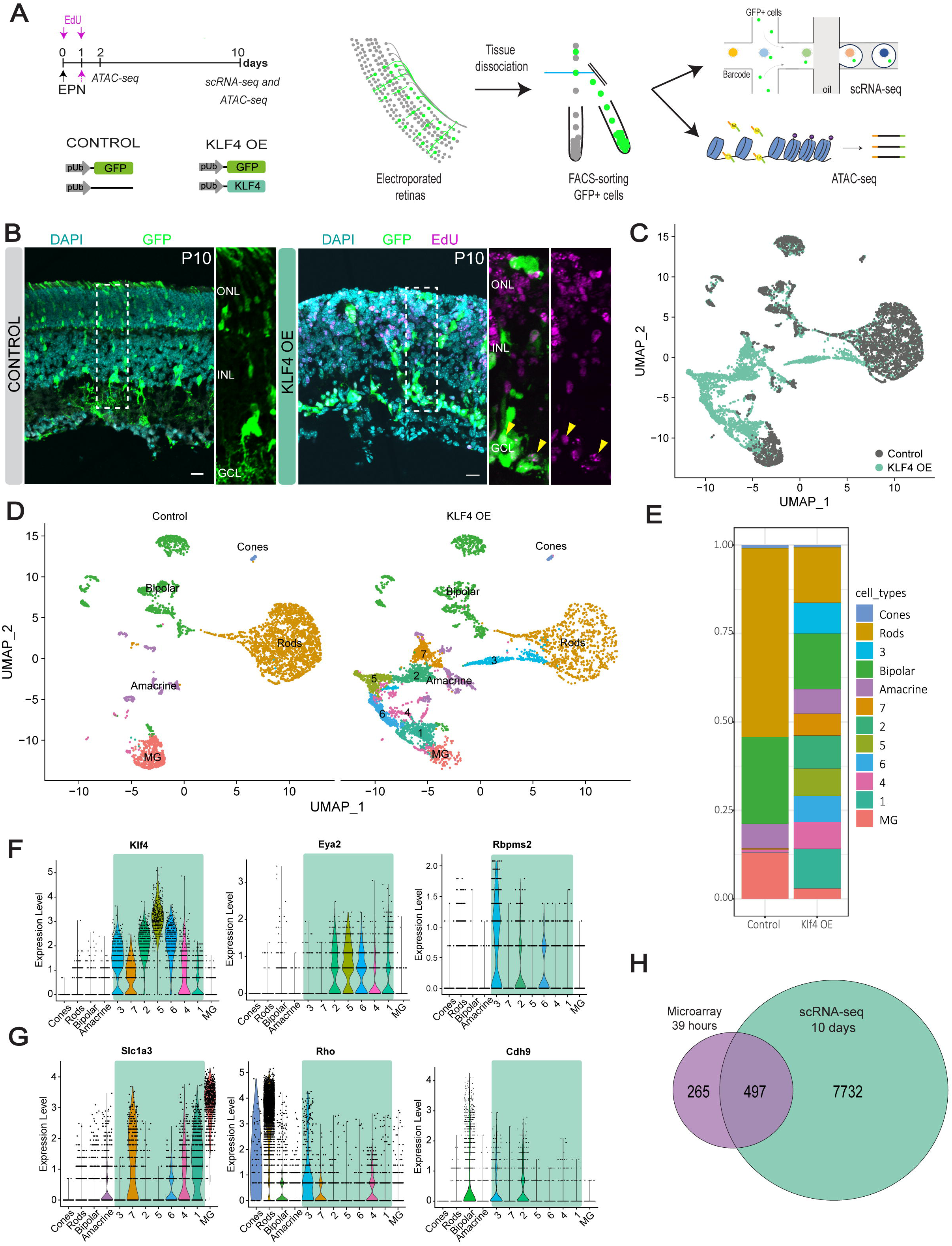
*Klf4* overexpression changes the fate of late retinal progenitors and induces heterogeneous transcriptional profiles. **(A)** Late retinal progenitors were electroporated with either pCST+pGFP (control) or pKLF4+pGFP (KLF4 OE, *Klf4* overexpression) constructs at postnatal day 0 (P0). Electroporated retinas were dissociated and GFP^+^ cells were isolated by FACS and processed for scRNA-seq or ATAC-seq. **(B)** Representative images of electroporated retinas show the laminar distribution of GFP^+^ cells in control and *Klf4* overexpression samples (n=4). Yellow arrowheads show GFP and EdU (magenta) double-positive cells. **(C)** UMAP visualization of scRNA-seq data reveals the distinct distribution of cells from the *Klf4* overexpression condition (turquoise) relative to control (grey). **(D)** Cluster analysis identified known retinal cell types and novel clusters enriched in the *Klf4*-overexpression condition, labeled 1-7. **(E)** Bar graph of the proportion of cells assigned to each cluster, split by experimental condition. **(F)** Violin plots show the expression of *Klf4* and genes previously reported to be upregulated 39 hours after *Klf4* overexpression (*Eya2* and *Rbpms2*) and **(G)** markers of late-born retinal cell types (*Slc1a3*, *Rho,* and *Cdh9*). Clusters induced by *Klf4* overexpression are indicated by a turquoise box. **(H)** Venn diagram of differentially expressed genes (DEGs) identified 39 hours (purple circle) or 10 days after *Klf4* overexpression (turquoise circle). ONL: outer nuclear layer; INL: inner nuclear layer; GCL: ganglion cell layer. Scale bar: 20µm.

Consistent with our previous report, 10 days after electroporation with *Klf4*, we observed the emergence of ectopic GFP^+^ cells in the inner retina, including within the ganglion cell layer (GCL), while control retinas had GFP^+^ cell bodies primarily localized to the ONL and inner nuclear layer (INL), and not the GCL. We also observed formation of retinal rosettes in regions densely electroporated with *Klf4* (**Figure 1B**). To ensure that the GFP^+^ cell distribution reflects genuine fate changes from late RPCs rather than potential artifacts arising from GFP transfer, we conducted lineage-tracing experiments. Specifically, cycling progenitors were labeled with EdU at P0 and P1, followed by *in vivo* electroporation at P1 (**Figure 1A**, magenta). Even though the two EdU pulses did not label all late-stage RPCs, we observed a subset of GFP^+^EdU^+^ cells at the GCL (**Figure 1B**, yellow arrowheads). These findings are consistent with our prior work in rat (Rocha-Martins et al., 2019), and confirm that *Klf4* overexpression (KLF4 OE) induces redistribution of cells to the GCL. This suggests that *Klf4* influences the fate of cell types generated from late RPCs.

To obtain transcriptome-wide expression profiles of the cell types induced by *Klf4* overexpression, we employed single-cell RNA sequencing (scRNA-seq). Late RPCs were electroporated on P0, followed by isolation of GFP-positive cells using fluorescence-activated cell sorting (FACS) 10 days after electroporation (**Figure 1A**). UMAP dimensional reduction revealed that *Klf4* overexpression in late retinal progenitors produced cells with largely distinct transcriptional profiles compared with controls (**Figure 1C, Table S1**). Unsupervised cell clustering analysis, followed by manual cell type annotation showed that the control sample was enriched for rod, Müller glia, bipolar, and amacrine cells, consistent with the expected late-born cell types (**Figure 1D, E**). Notably, *Klf4* overexpression produced new clusters that we designated as 1 to 7, and decreased proportions of cells in rod, bipolar, and Müller glia clusters, suggestive of changes in cell fate (**Figure 1D and E**). We observed higher *Klf4* levels in the new clusters, along with the expression of RGC genes (**Figure 1F, Table S2**). These KLF4-induced cells, however, retain some expression of genes associated with later born cell types, suggesting a mixed identity (**Figure 1G, Table S2**).

Previously, we had assessed gene expression changes 39 hours after *Klf4* overexpression using microarray profiling and identified 762 differentially expressed genes (DEGs) when compared to control GFP^+^ cells (Rocha-Martins et al., 2019). Here, we detected 8,229 DEGs between control and KLF4-induced cells in our scRNA-seq dataset. We compared these two gene lists to determine what early transcriptional changes are sustained as cells differentiate and mature. The intersection of these datasets contained 497 genes that were commonly regulated at both time points (**Figure 1H** and **Table S3**). Further examination of their expression dynamics showed that the majority displayed consistent regulation, with those induced at early stages remaining upregulated at 10 days (**Table S3**).

### Chromatin accessibility is widely altered with Klf4 overexpression

*Klf4* is a well-established pioneer transcription factor (Soufi et al., 2015), suggesting it likely remodels chromatin accessibility to regulate retinal cell fate. To identify genome-wide alterations in chromatin accessibility following *Klf4* overexpression, we performed ATAC-seq analysis relative to control conditions. As with our scRNA-seq experiment, GFP and KLF4 vectors were electroporated into late progenitors at P0, and GFP^+^ cells were collected for ATAC-seq after 2 and 10 days. We identified a total of 149,519 genome-wide chromatin regions that were open in at least two samples. Through differential accessibility analysis, using the number of cut sites within these peaks, we identified 5,744 genomic regions that were more open in response to *Klf4* overexpression after 10 days, and 2,260 regions that were less accessible (**Figure 2A, Table S4**). By visual inspection, we observed that the KLF4 induced open chromatin regions were progressively closed in control samples from P2 to P10, indicating that these regions are developmentally regulated. These KLF4 open regions already showed greater accessibility than control 2 days after *Klf4* overexpression, with accessibility further increasing after 10 days, suggesting that KLF4 induced changes in the chromatin landscape rapidly and sustainably (**Figure 2A**). Motif enrichment analysis revealed that several KLF family motifs were enriched in regions that became accessible following *Klf4* overexpression (**Figure 2B, Table S5**).

**Figure 2.**
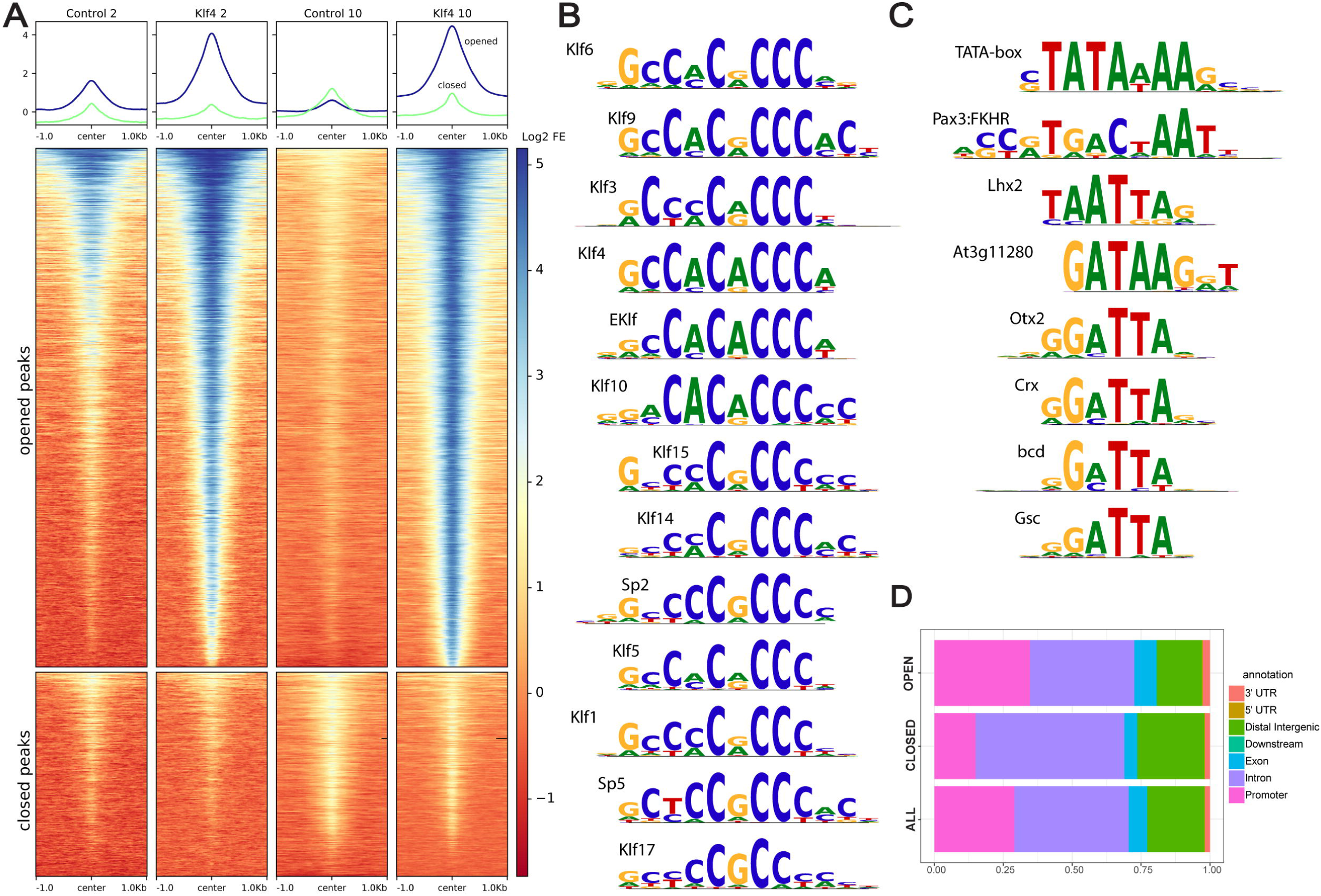
Chromatin accessibility changes following *Klf4* overexpression. ATAC-seq performed after two or ten days of *Klf4* overexpression identified regions exhibiting increased (opened) or decreased (closed) chromatin accessibility. **(A)** Heatmap of chromatin landscape surrounding differentially accessible regions across experimental conditions. Blue indicates enrichment of transposase cut sites, and red indicates depletion of cut sites. **(B)** Top 15 transcription factor motifs enriched in regions that gained chromatin accessibility following *Klf4* overexpression. **(C)** Selected motifs enriched in regions that lost chromatin accessibility upon *Klf4* overexpression. **(D)** Bar plot of the genomic annotation peaks according to their location relative to nearest gene, for opened, closed and all peaks.

Those peaks that showed reduced accessibility 10 days after *Klf4* overexpression relative to control, appear to be developmentally regulated, normally progressively opening from P2 to P10. While these peaks still opened from 2 to 10 days after *Klf4* overexpression, their accessibility appears reduced relative to control at both timepoints (**Figure 2A**). Motif analysis of closed regions at 10 days did not identify enrichment of any KLF family motifs. Rather, these regions were enriched for CRX, OTX2, and LHX factor motifs, which play a role in the specification and differentiation of late cell types (Zuber et al., 2003) (**Figure 2C, Table S6**).

To identify which genes are likely regulated by accessibility of these regions, we annotated all peaks for their position relative to the nearest gene. Overall, KLF4-induced open regions, when compared to all accessible peaks, were slightly less likely to reside in distal intergenic regions, and more likely to be found in promoter regions, which serve as the core regions that facilitate gene regulation (Riethoven, 2010) (**Figure 2D**). KLF4 closed peaks were more likely to be found in introns than in promoter regions when compared to all accessible regions (**Figure 2D**).

We examined whether the changes observed in chromatin accessibility were likely to predict gene expression changes by intersecting the differentially regulated peaks with the list of DEGs from scRNA-seq. Our findings revealed that opened regions are more likely to be located near upregulated genes, while closed regions are more likely to be near downregulated genes (**Figure 3A**). 39.6% of opened ATAC-seq regions are proximal to genes up-regulated in scRNA-seq, indicating a strong association. This pattern reinforces the idea that KLF4-induced changes in chromatin accessibility directly control transcriptional expression. To evaluate whether KLF4 might directly regulate specific genes, we identified all ATAC-seq peaks containing a KLF4 binding motif. We identified 1,262 peaks that contain KLF4 motifs, are more accessible after *Klf4* overexpression, and are proximal to genes upregulated after *Klf4* overexpression. These regions could indicate a direct effect of KLF4 (**Figure 3B**). For example, this pattern can be seen for *Pax6*, which has one significantly opened chromatin region that contains a KLF4 motif (red asterisk), and is significantly upregulated after *Klf4* overexpression (**Figure 3C, Table S1**). In addition, we identified 122 peaks in open regions, containing a KLF4 motif but that are proximal to downregulated genes, suggesting that KLF4 may act as a repressor of these genes. An example of this potential relationship is at the *Casz1* locus, where three sites have significantly increased chromatin accessibility and two of these contain KLF4 motifs (red asterisk), while *Casz1* is downregulated in our scRNA-seq analysis (**Figure 3C, Table S1**).

**Figure 3.**
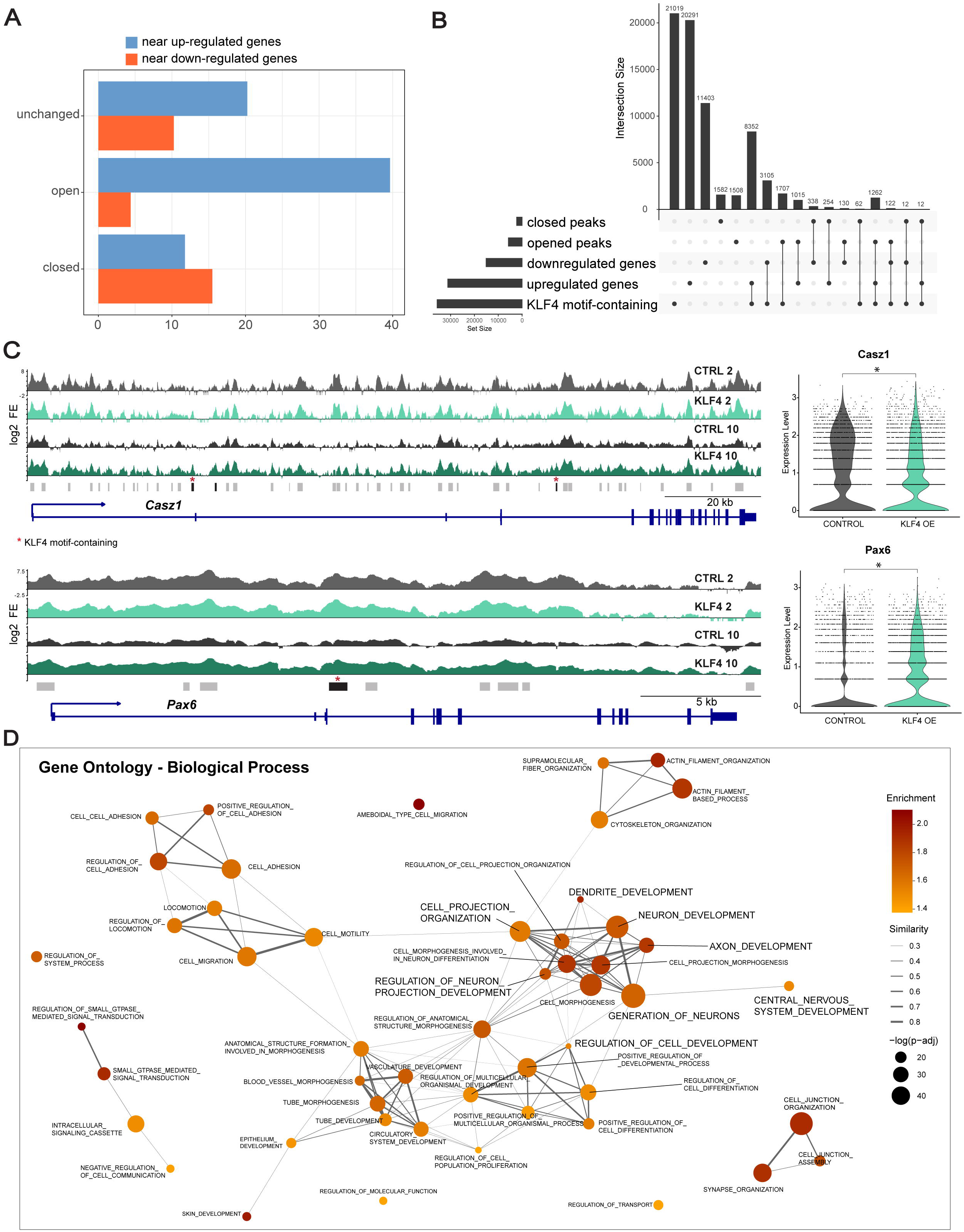
Changes in chromatin accessibility induced by *Klf4* overexpression correlate with transcriptional alterations. **(A)** Bar graph showing the proportion of opened, closed, or unchanged ATAC-seq peaks annotated for up- or down-regulated genes from scRNA-seq. **(B)** UpSet plot demonstrates the intersection between closed and opened peak annotations, up-and downregulated genes identified by scRNA-seq, and genomic regions containing KLF4 motifs. **(C)** Data for representative genes *Casz1* and *Pax6*, both containing regions of increased chromatin accessibility and decreased or increased gene expression, respectively. Genome tracks show ATAC-seq results averaged by experimental condition. Bars beneath the log2 fold-enrichment tracks indicate all peak regions. Those peaks significantly opened in *Klf4* overexpression are shown in black, and peaks containing a KLF4-binding motif are marked with a red asterisk. Violin plots on the right display gene expression levels derived from the scRNA-seq data, stratified by condition. **(D)** Gene ontology analysis of genes associated with opened peaks. The 50 biological process pathways with the lowest p-value are represented as dots, with dot size proportional to the corrected p-value and color indicating fold enrichment. The Jaccard index of overlapping genes denotes the similarity between pathways by connecting line thickness. Similarities below 0.25 are not shown.

We conducted gene ontology analysis for genes proximal to regions that became more accessible (open) after *Klf4* overexpression to identify which pathways are affected by KLF4-induced chromatin landscape changes. The significantly enriched biological processes were consistent with the observed phenotype, including neuron generation, neuron development, regulation of cell differentiation, and cell projection organization (**Figure 3D**).

### KLF4 overexpression results in cells with RGC-gene expression

Previously, we demonstrated that *Klf4* overexpression initiates an RGC developmental program and induces RGC morphological features in late retinal progenitors. This was evident by the position of the electroporated cells in the ganglion cell layer and their extension of projections toward the optic nerve head (Rocha-Martins et al., 2019) (**Figure 1B**). To gain further insight into the identity of KLF4-induced cells, we compared our scRNA-seq data with a published dataset of all retinal cell types across development from embryonic day 11 (E11) to P14 (Clark et al., 2019).

Integration of these two datasets resulted in a clear coincidence of late cell types (rods, bipolar cells, and Müller glia) by UMAP reduction, demonstrating transcriptional resemblances and accurate annotation (**Figure 4A, B**). KLF4-induced clusters similarly colocalized with or were near established retinal cell types, including RGCs, suggesting transcriptional similarities (**Figure 4A and B**). Automated annotation using the same retina development dataset as a reference likewise reinforced our control cell identities and annotated our novel populations as RGCs, amacrine cells, and others (**Figure 4C**). Specifically, we observed that clusters 2 and 3 and some cells from clusters 5 and 7 colocalized with developmental RGCs (zoomed in **Figure 4D**). Given the observation that cells acquire an RGC-like transcriptomic profile, we examined the expression of 75 enriched or specific RGC genes identified through analysis of the retina development dataset (Clark et al., 2019) (**Figure 4E**, green box). Notably, several of these RGC-associated genes were expressed within specific KLF4-induced clusters, particularly those adjacent to the bona fide RGC cluster (**Figure 4D**). Altogether these findings substantiate the observation that *Klf4* overexpression redirects the fate of retinal cells generated from late RPCs including inducing RGC-like cells.

**Figure 4:**
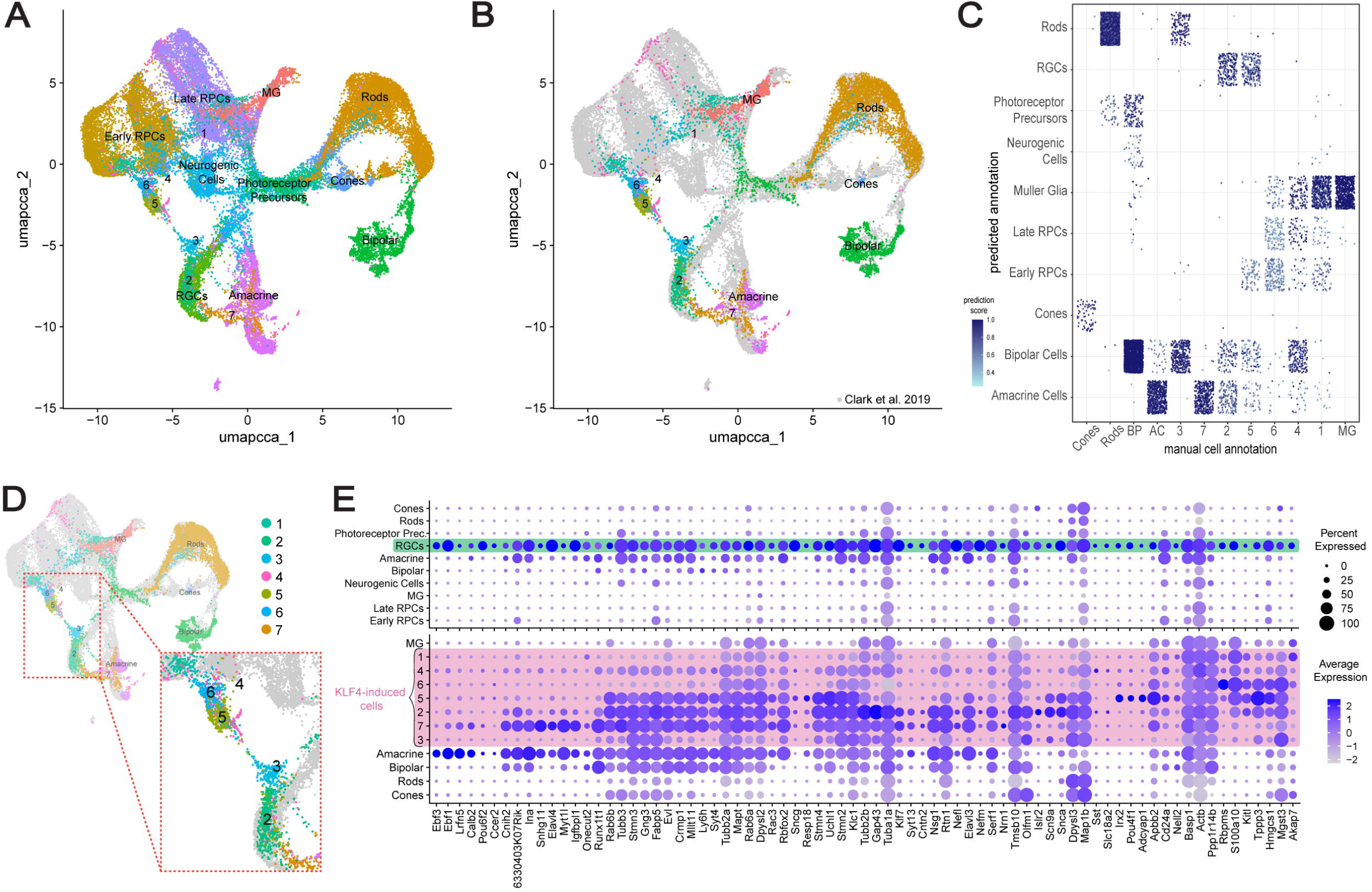
KLF4-induced cells exhibit molecular similarities to embryonically generated retinal ganglion cells. **(A)** UMAP-based dimensional reduction of the *Klf4*-overexpression scRNA-seq dataset integrated with a reference dataset of retinal development dataset demonstrates that clusters emerging under the *Klf4* overexpression condition map closely to retinal ganglion cell (RGC) populations. **(B)** UMAP visualization with reference retinal cells shown in grey. **(C)** Automated cell-type annotation compared with manual cell annotation predicts a subset of cells from clusters 2 and 5 as RGCs. Cell type prediction scores are represented by a blue gradient, with darker shades indicating higher confidence. **(D)** Magnified view of the indicated UMAP region (red dashed box) highlights cells positioned in proximity to endogenous RGCs. **(E)** Dot plot depicting the expression of 75 genes enriched in RGCs across reference retinal cell types and *Klf4*-overexpression scRNA-seq clusters.

### KLF4 induces an RGC program in late retinal progenitors

We found several genes implicated in RGC neurogenesis to be upregulated following *Klf4* overexpression (Rocha-Martins et al., 2019) including *Atoh7* and *Eya2* which have roles in RGC competence and specification (Lyu & Mu, 2021). Additionally, we detected the expression of RGC genes such as *Pou4f2*, *Pou4f1, Foxp1*, *Stmn2*, *Sox11,* and *Gap43* predominantly in the newly formed clusters (1-7) induced by *Klf4* overexpression (**Figure 5A and B**). Notably, these genes did not show uniform expression across every new cluster, suggesting that KLF4 only partially initiates the RGC transcriptional program.

**Figure 5:**
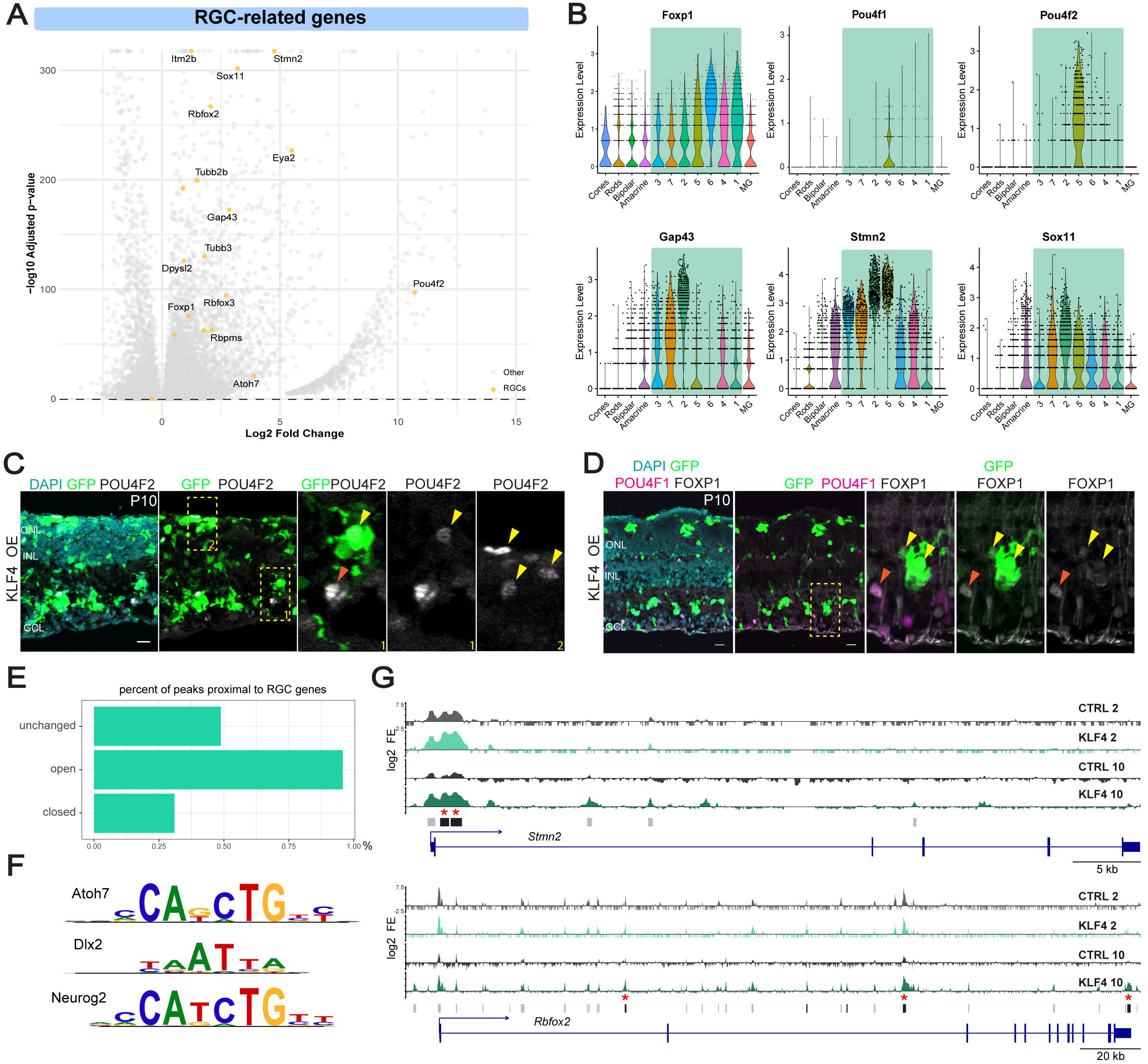
KLF4 overexpression activates RGC-associated genes and remodels their chromatin accessibility. **(A)** Volcano plot showing differentially expressed genes (DEGs) identified by scRNA-seq following Klf4 overexpression with genes implicated in RGC development labeled. KLF4 up-regulated genes display positive log2 fold change values. **(B)** Violin plots depict the expression patterns of selected RGC-associated genes across clusters. **(C)** Immunofluorescence staining for POU4F2 (grey), GFP (green), and DAPI (blue), and **(D)** for POU4F1 (magenta) and FOXP1 (grey) validate the scRNA-seq data (representative images: n=2 for POU4F2 and n=4 for FOXP1). Yellow arrowheads indicate *Klf4* overexpression cells expressing RGC genes (POU4F2^+^GFP^+^ or FOXP1^+^GFP^+^); orange arrowheads mark endogenous RGCs (POU4F2^+^GFP^-^ or FOXP1^+^POU4F1^+^GFP^-^). **(E)** Bar plot of the percentage of ATAC-seq peaks proximal to RGC genes. **(F)** RGC-related transcription factor binding motifs are enriched within regions that gained chromatin accessibility. **(G)** Genome browser tracks across the *Stmn2* and *Rbfox2* loci demonstrate increased chromatin accessibility in the *Klf4* overexpression condition (turquoise tracks) relative to controls (grey tracks). Significantly opened peaks are colored in black, and peaks containing a KLF4 motif are marked with red asterisks. ONL: outer nuclear layer; INL: inner nuclear layer; GCL: ganglion cell layer. Scale bar: 20µm.

We analyzed the expression of *Pou4f1, Pou4f2* and *Foxp1* by immunostaining retinas following *Klf4* overexpression. We observed POU4F2^+^GFP^+^ cells both ectopically positioned in the ONL (**Figure 5C-2**) and appropriately located in the GCL (**Figure 5C-1**). Additionally, FOXP1^+^GFP^+^ cells were observed following *Klf4* overexpression (**Figure 5D**, gray). In contrast, although *Pou4f1* was expressed in cluster 5 (**Figure 5B**), POU4F1^+^GFP^+^ cells were not detected through immunostaining which may suggest it is a rare population or that it presents low induction levels (**Figure 5D**, magenta). These immunostaining results reinforce our observation that *Klf4* overexpression induces some but not all RGC-related genes in electroporated cells.

*Klf4* overexpression not only induced expression of RGC-related genes, but also preferentially opened regions near RGC genes, as shown by our ATAC-seq data (**Figure 5E**). 0.96% of opened ATAC-seq regions are near RGC-related genes, compared with 0.49% of unchanged regions, consistent with the up-regulation of the RGC transcriptional program we described. This supports an intrinsic role in recognizing genomic regions to induce the RGC program. Binding motifs of critical RGC-promoting transcription factors, including ATOH7, DLX2, and NEUROG2, were enriched in KLF4-opened regions, allowing for the possibility of indirect or cooperative regulation of the RGC developmental program (**Figure 5F**). We identified specific peaks in upregulated RGC genes, including *Stmn2* and *Rbfox2*, loci that have increased accessibility after *Klf4* overexpression and contain KLF4 motif sites (red asterisks), supporting the possibility of direct gene regulation (**Figure 5G**). Collectively, these findings demonstrate that KLF4 can redirect the fate of late RPCs, enabling the emergence of RGC-like cells by the activation of a cell-type-specific program. However, these reprogrammed cells do not fully mature into authentic RGCs, indicating incomplete differentiation.

### Axonogenesis-related genes are differentially regulated in KLF4-induced cells

Our previous findings demonstrated that the RGC-like cells generated following *Klf4* overexpression can extend axonal projections toward the optic nerve head, however, these axons fail to progress beyond this point in the visual pathway (Rocha-Martins et al., 2019). To understand this limitation on axon projection we examined RGC axonogenesis genes. Gene ontology (**Figure 3D**) and analysis of gene expression profiles revealed that genes related to axonogenesis, and axonal regeneration such as *Pou4f2* (**Figure 5A**)*, Sema3b, Tppp3, Robo2,* and *Elf1* (Nakaya et al., 2012; Oliveira-Valença et al., 2025; Thompson et al., 2009) were upregulated after *Klf4* overexpression (**Figure 6A and C**). Conversely, we observed the induction of genes known to inhibit axonogenesis or regenerative capacity including *Klf13*, and *Socs3* (Moore et al., 2009; Smith et al., 2009), which may constrain the pro-axonogenesis program (**Figure 6A and C**).

**Figure 6:**
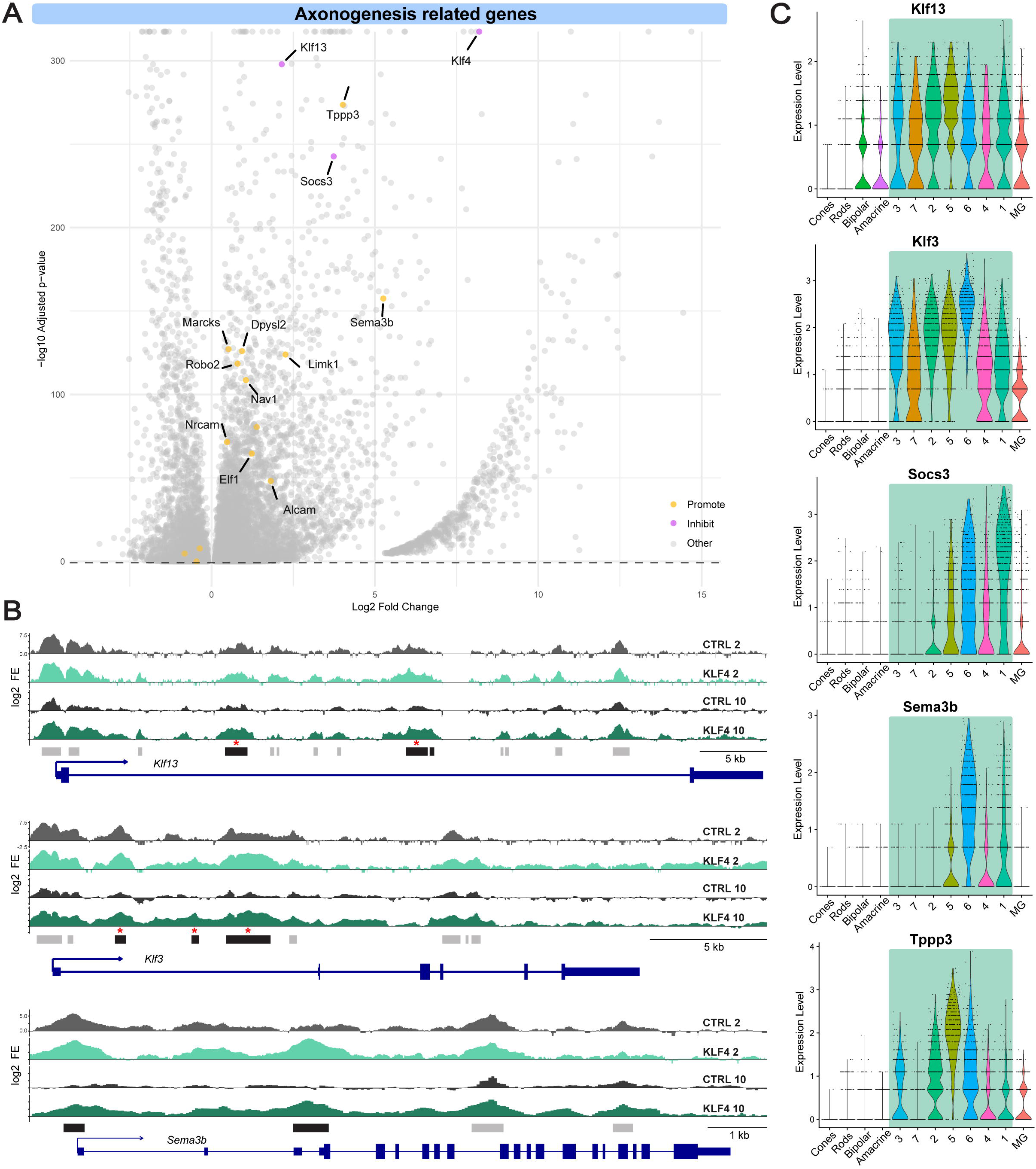
Genes regulated by KLF4 may underlie the limited axonal growth in RGC-like cells. **(A)** Volcano plot of all *Klf4* overexpression DEGs identified by scRNA-seq. Genes associated with axonogenesis are labeled with yellow or purple dots, indicating those that promote or inhibit axonal growth, respectively. **(B)** ATAC-seq genome browser tracks across the *Klf13*, *Klf3*, and *Sema3b* loci reveal increased chromatin accessibility in the *Klf4* overexpression condition (turquoise tracks) relative to the controls (grey tracks). Red asterisks mark KLF4 motif-containing peaks and significantly opened peaks are shown in black. **(C)** Violin plots display the expression patterns of selected axonogenesis-related genes across cell clusters. KLF4-induced clusters are highlighted by a turquoise box.

Prompted by the observed induction of *Klf13*, we investigated whether other KLF factors were similarly affected. Indeed, we observed changes in chromatin accessibility in several KLF factor loci, including *Klf3* (**Figure 6B**). In addition, the scRNA-seq revealed upregulation of *Klf3*, *Klf10*, *Klf8*, *Klf7,* and *Klf12* after *Klf4* overexpression (**Table S1**). These results show that *Klf4* overexpression can induce genes involved in axonogenesis, however, this program is insufficient to overcome inhibition of axon projection, potentially by the contribution of other KLF family members (Moore et al., 2009).

## DISCUSSION

Here, using single cell RNA sequencing, we demonstrate that *Klf4* overexpression in late RPCs redirects their cell fate, inducing expression of RGC-related genes and acquisition of RGC-like cell identity. ATAC-seq revealed that these changes in gene expression are reflected in genome-wide changes in chromatin accessibility, consistent with the established role of KLF4 as a pioneer transcription factor. We found upregulation of genes related to RGC axonogenesis, but our prior work showed that KLF4-induced cells do not extend axons beyond the initial segment of the optic nerve (Rocha-Martins et al., 2019). Genes known to inhibit axon growth and genes normally expressed in other retinal neurons are also expressed in these cells, likely preventing them from adopting a full RGC identity and morphology.

The current study confirmed our previous data across species (Rocha-Martins et al., 2019) and demonstrated that *Klf4* overexpression indeed directs late RPCs to generate cells with new expression profiles including RGC-like cells which reach the inner retina. Furthermore, we investigated the potential mechanisms by which KLF4 influences this change in cell fate. The identity of KLF4-induced cells resembles RGCs generated during the embryonic window, as revealed by integration of our scRNA-seq dataset with a developing retina reference (Clark et al., 2019). These RGC-like cells are not transcriptionally identical to embryonically derived RGCs, reinforcing what we have shown by EdU lineage tracing: that the KLF4-induced cells are derived from dividing late RPCs rather than an artifact of GFP transfer between cells. Some of the new clusters identified by scRNA-seq retain partial expression of Müller glia and rod genes, and do not express the full RGC regulatory network. This heterogeneity might be due to variations in the copy number or levels of *Klf4* overexpression, but also to competence restrictions or intrinsic variability of late RPCs (Trimarchi et al., 2008).

A key property of RGCs is the projection of axons to central targets. We observed changes in gene expression of several factors that either favor or inhibit axonogenesis, which may explain our prior observation of induced cells projecting to the optic nerve head but not to the brain (Rocha-Martins et al., 2019). *Klf4* itself and *Klf13* are both highly upregulated and are known to inhibit this process (Moore et al., 2011). Other KLF family genes were also regulated, including *Klf3*, *Klf10*, and *Klf7*, with the last being the only one known to promote axonogenesis. Moore et al. (2011) showed that inhibition is prevalent even when combining KLF factors with opposing effects, reinforcing the idea that KLF4 and KLF13 may act to repress axon growth despite the presence of other genes that promote it.

Since KLF4 induced an RGC-related gene expression program, we sought to determine whether the observed gene expression changes were associated with chromatin reorganization. Our findings show that after 10 days of *Klf4* overexpression there are genome-wide chromatin accessibility changes, primarily increased accessibility within regions enriched with KLF motifs. This suggests that this is a direct effect of KLF4 binding to, and opening, chromatin. The opened chromatin was also enriched for motifs of factors related to RGC genesis, such as ATOH7, DLX2, DLX1, NEUROD1, ISL1, POU4F3, FOXP1, and NEUROG2.Increased access for these transcription factors may provide a clue as to how the RGC transcriptional program was broadly altered (Lyu & Mu, 2021). This suggests a role for KLF4 as a pioneer factor capable of opening cell-type specific regulatory regions (Soufi et al., 2015). Chromatin regions with significantly decreased accessibility were enriched for motifs that are associated with late cell-type specification, including OTX2, CRX, and LHX2. We hypothesize that these conformational changes are due to upregulation in the RGC transcriptional program and both indirect and direct effects of *Klf4* overexpression, but we cannot rule out the possibility that the reduced accessibility for late cell-type genes may be a consequence of the reduction in the number of cells with this identity, since we performed a bulk analysis.

We also observed that KLF4 induces changes in late RPCs 39/48 hours after its overexpression, which persists for 10 days. That was observed not only at the transcriptional level but also at the epigenetic level. This observation shows that KLF4 can promote long-lasting changes in cells and influence their cell fate. Alternatively, cells may be transcriptionally frozen and cannot fully differentiate into a specific cell identity.

To identify potential direct transcriptional targets of KLF4, we analyzed open ATAC-seq regions proximal to differentially expressed genes and containing KLF4 binding motifs. We described several candidates for direct upregulation by KLF4 with established roles in RGC genesis, including *Eya2*, *Atoh7*, *Rbfox2*, and *Stmn2* (Dvoriantchikova et al., 2015; Gao et al., 2014; Gu et al., 2020; Laboissonniere et al., 2019). The sustained expression of *Atoh7* in KLF4-induced cells contrasts with its developmental expression pattern, during which *Atoh7* is transiently expressed (Brzezinski et al., 2012). We also found upregulation of *Notch1*, shown to inhibit photoreceptor differentiation (Chen & Emerson, 2021; Jadhav et al., 2006; Yaron et al., 2006), which could contribute to the shift in cell fate.

A direct repressive role for KLF4 has been previously described (Kaczynski et al., 2003) and, indeed, we found several downregulated genes as candidates for direct KLF4 regulation. One of the genes is *Casz1*, which is known to be a competence factor present in mid-late progenitors during retinal development (Mattar et al., 2015). Notably, among the genes that are significantly upregulated at 39 hours after *Klf4* overexpression, most remain upregulated after 10 days. Similarly, changes in chromatin accessibility that are significant at 10 days are already visible at 48 hours. Together, this suggests that KLF4-induced changes are both rapid and long-lived.

Altogether, we show that *Klf4* overexpression in late RPCs can broadly induce chromatin remodeling and drive changes in gene expression, including activation of an RGC-related gene expression program. These findings support KLF4, potentially in combination with other factors, as a promising candidate for the development of regenerative protocols targeting Müller glia, which share transcriptional and epigenetic similarities with late RPCs, and serve as a likely retinal neuron regenerative source (Bernardos et al., 2007; Blackshaw et al., 2004; Ooto et al., 2004; Roesch et al., 2008).

## Author contributions

Conceptualization: M.S.S., V.M.O-V ; Data curation: V.M.O.-V., J.M.R., F.C., A.B., M.L.V., M.S.S.; Formal analysis: V.M.O.-V., J.M.R., M.L.V., M.S.S.; Funding acquisition: A.B., M.L.V., M.S.S.; Investigation: V.M.O.-V., J.M.R., F.C.; Methodology: V.M.O.-V., J.M.R.; Resources: M.L.V., M.S.S.; Supervision: M.S.S.; Validation: V.M.O.-V.; Visualization: M.S.S., V.M.O-V; Writing – original draft: V.M.O.-V., M.S.S.; Writing – review & editing: V.M.O.-V., J.M.R., A.B., M.L.V., M.S.S.

## Data availability

Raw and processed high-throughput sequencing data generated in this study have been deposited in GEO under accession numbers GSE329012 and GSE329113.

---

**Table S1: Differentially expressed genes (*Klf4* overexpression vs. Control).**

**Table S2: Set of genes expressed in each cluster from scRNA-seq data.**

**Table S3: Intersection between *Klf4* overexpression scRNA-seq and microarray dataset.**

**Table S4: Count peaks annotation for close and open regions.**

**Table S5: Motif annotation for opened regions after *Klf4* overexpression (10 days).**

**Table S6: Motif annotation for closed regions after *Klf4* overexpression (10 days).**

**Table S7: Gene ontology analysis based on chromatin accessibility.**

